# Precipitation frequency and predictability interactively affect lizard life-history traits in absence of water shortage

**DOI:** 10.64898/2026.01.26.701707

**Authors:** Rebeca Vicente Moreno, Patrick S. Fitze

## Abstract

Current climate change leads to longer frequencies and reduced predictability of climatic parameters. Recent studies have highlighted the importance of considering multiple environmental factors, but experimental evidence on how species respond to their combined effect remains scarce. Here, we experimentally manipulated precipitation frequency and predictability and tested how they affect body size, growth, and survival using the common lizard (*Zootoca vivipara*) as a model species. Longer precipitation frequency negatively affected adult growth and male survival. Predictability influenced body size-dependent survival of yearlings and adults in certain frequency treatments. In yearlings, treatment-induced growth differences compensated for treatment-induced differences in size-dependent survival, resulting in no size differences during reproduction. In adults, treatment-induced differences in size-dependent survival were not compensated for, resulting in body size differences during reproduction among treatments. Consequently, precipitation frequency and predictability had a joint effect on life-history traits. Our results demonstrate that, even without water shortage, small differences in the frequency and predictability of precipitation affect population demography and life-history traits. This indicates that integrating the interactive action of different climatic parameters will be key to understanding and better anticipating future impacts of climate change on species.

## Introduction

Understanding how organisms respond to changes in their environments is a fundamental theme in ecology, especially in the context of global climate change. Annual mean temperature, total precipitation, and changes therein are among the most frequently examined climatic parameters^1^.

However, the current climate change not only impacts average environmental conditions, but also amplifies variability, reduces predictability, and changes the frequency of environmental parameters^2–4^.

Recent research shows that reduced environmental predictability influences life-history traits^5^, and has negative consequences for population dynamics^6^. Other studies showed that less predictable precipitation produces relatively small effects on adult survival and reproduction^7^, affects elaborate ornamentation^8^, and negatively affects survival, growth, and body condition of competitively inferior age classes (yearlings and juveniles^6^). This suggests that different age-classes exhibit different sensitivities to environmental predictability^7,8^. However, age class dependent effects may also be explained by life-history theory and the trade-off between current and future reproduction^9^. Different ages may show different resolutions of the trade-off between survival and growth depending on the environmental predictability, and these resolutions may be sex-specific^10,11^. In this case, population dynamics may not be influenced by environmental predictability, contrasting with studies showing that even small, non-significant survival differences can have major consequences for population growth rate, and suggesting that individuals may not be able to cope with reduced environmental predictability^6^. These experimental findings support the theoretical predictions that environments with lower predictability may lead to increased population extinction^12^.

In many regions of the world, global climatic change does not change the annual precipitation amount, but it strongly affects alterations in precipitation frequency, or, in other words, the length of the recurrence interval between two precipitation events^2^. A scenario characterized by fewer but larger precipitation events automatically results in longer periods without rainfall, leading to periods of low soil moisture, particularly in the upper soil layers^13,14^. Extended drier periods also lead to soil sealing, which hinders precipitation from penetrating into deeper soil layers and enhances the risk of soil drying^15^. Low water availability is known to negatively affect many terrestrial species, including the hydrophilic *Zootoca vivipara*^16^.

Global change imposes additive and interactive changes in multiple environmental factors. Interactive effects seem to be more common than additive effects^17^, and synergistic interactions (i.e., when the combined effects are greater than the sum of the individual effects) can entail a higher risk of population decrease^18,19^. However, interactive effects may also moderate the effects of the single factors (antagonistic interaction)^20^. Therefore, single-factor experiments may not reveal how global change affects population dynamics, and only simultaneous testing of multiple parameters on life-history traits and population demography will provide a more precise understanding^17^.

Here, we experimentally investigated whether and how two precipitation parameters, altered by climate change, affect demography and life-history traits, using the common lizard (*Zootoca vivipara*) as a model species. We manipulated precipitation frequency and intrinsic precipitation predictability (used interchangeably with precipitation predictability in this article) to which lizards were exposed using a crossed 3 x 2 design and two semi-natural populations per treatment combination, and repeated the experiment over two consecutive experimental years (experimental year refers to a period between June and May of the following year). All lizard populations consisted of the same standardized population structure (i.e., age class and sex ratio), and after one year, lizards were recaptured to quantify treatment effects on body size, growth rates, and annual survival. These life-history traits are important determinants of population dynamics^21^, given that they are closely linked to population growth rate^22^. Body size predicts maturity^23–25^ and thereby influences recruitment to the population, fecundity, and ultimately fitness^26,27^. Moreover, differences in body size may arise from variations in growth rates^7^ and/or size-dependent survival^28^, due to a different resolution of the trade-off between survival and growth or survival and reproduction^10^. Conversely, the lack of effects on body size may be the consequence of compensatory plastic responses^21,27,29^. Elucidating the involved mechanisms is thus crucial for the understanding of the treatment consequences. According to life-history theory, the allocation of resources among growth, reproduction, and survival is shaped by inherent constraints and trade-offs, with optimal allocation patterns depending on relative age-and sex-specific fitness sensitivities^10,11^. In less predictable and potentially more stressful environments, we expected reduced growth rates and smaller adult body size, particularly among juveniles, as well as lower survival in age classes less able to buffer environmental changes. We therefore predicted (1) significant negative effects of less frequent and less predictable precipitation on growth, body size, and survival, and (2) that lizards may exhibit compensatory mechanisms to mitigate the negative treatment effects. We also tested (3) if differences in body size arose due to differential growth or alternatively, (4) due to body size-dependent survival selection.

## Results

### Adults

In adults, survival was affected by a significant interaction between FREQ treatment, sex, and year (*χ^2^*_2_ = 6.665, *P* = 0.036), and there was no significant effect of PRED treatment (*χ^2^*_1_ = 0.115, *P* = 0.734). Post-hoc analyses showed that in females of the 2019-20 experiment, survival was significantly lower in F24 than in F48 (*P* = 0.034; Fig. 1a) in 2019-20. In males, survival was significantly lower in F96 compared to F24 and F48 (*P* = 0.039 and *P* = 0.018, respectively; Fig. 1a), and no other post-hoc comparisons were significant (all other *P* > 0.05). Body size at recapture was significantly influenced by an interaction between FREQ, PRED, and sex (*χ^2^*_2_ = 11.782, *P* = 0.003). Post-hoc analyses revealed that in females, body size in LP was significantly bigger than in MP in F24 (*P* = 0.032; Fig. 1b), and in F48, MP was significantly bigger than LP (*P* = 0.043; Fig. 1b), whereas no significant effects were observed in F96 (*P* = 0.250). In males, body size in F24 was significantly bigger in MP than in LP (*P* = 0.031), and no significant differences existed in the other FREQ treatments (all *P* > 0.2). Within PRED treatment levels, female body size was bigger in F24 than in F48 in the LP treatment (*P* = 0.047; supplementary Fig. S1), and in the MP treatment, it was bigger in F48 than in F24 and F96 (*P* = 0.042 and *P* = 0.005, respectively; supplementary Fig. S1). In males, no significant contrasts existed among FREQ treatments (all *P* > 0.1). Since by experimental design, the average and variance of SVL at the start of the experiment (SVL_Initial_) were similar across all treatments and enclosures (all *P* > 0.8), any differences in body size observed at the end of the experiment must be due to treatment differences in growth rates, and/or treatment differences in size-dependent survival. Growth rate significantly differed among FREQ treatments (*χ^2^*_2_ = 6.751, *P* = 0.034). Lizards grew significantly faster in F48 compared to F96 (*P* = 0.01; Fig. 2a). SVL_Initial_ of released adult lizards significantly differed among survivors and non-survivors, in interaction with FREQ, PRED, and sex (*χ^2^*_2_ = 11.434, *P* = 0.003). In females exposed to F24 and the MP treatment, surviving females had significantly smaller initial SVL compared to females that died over the course of the experiment (*P* = 0.026; Fig. 2b). In males exposed to F24 and the MP treatment, surviving males had significantly bigger initial SVL than those who died over the course of the experiment (*P* = 0.046; Fig. 2b), and no other contrast was statistically significant (all *P* > 0.3). The PRED effect on final body size of adult females and males in F24 (Fig. 1b) did thus arise due to body size-dependent mortality (Fig. 2b; also see Appendix S1, supplementary Fig. S2 for a visual representation of the causes of the observed effects on adult SVL_Final_).

**Figure 1.**
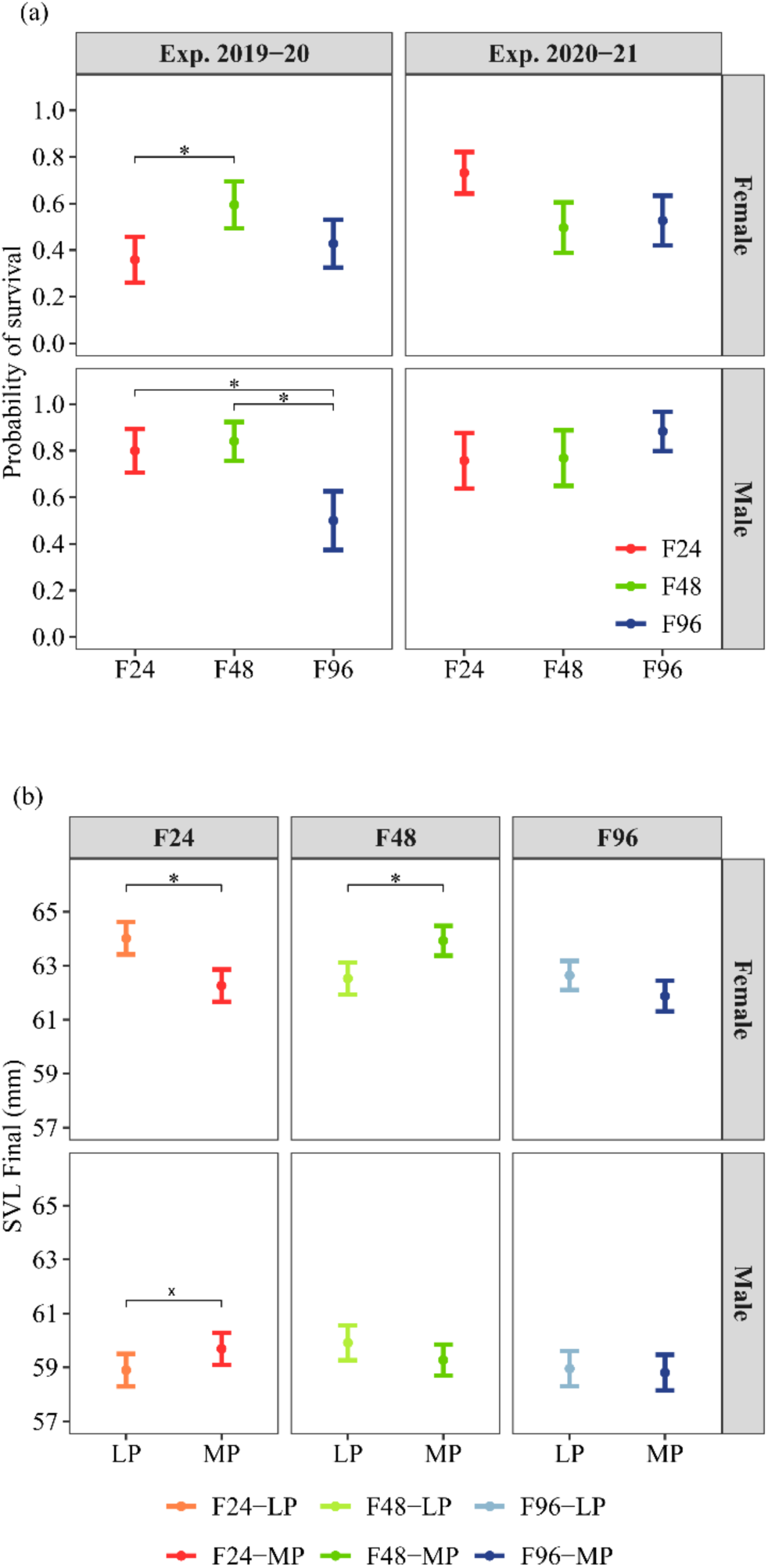
Treatment effects on adult survival (**a**) and adult body size at recapture (**b**). Predicted means ± SE per FREQ treatment (Frequency treatment: F24: 24-hour recurrence interval; F48: 48-hour interval; F96: 96-hour interval), experimental year (Exp.) and sex are shown in panel (**a**), and the interaction of FREQ, PRED (Predictability treatment: MP, more predictable; LP, less predictable) and sex in panel (**b**). Horizontal lines within the graphs indicate significant post-hoc contrasts: * *P* < 0.05; ** *P* < 0.01; *** *P* < 0.001; ^x^ P < 0.05 when analyzing males alone.

**Figure 2.**
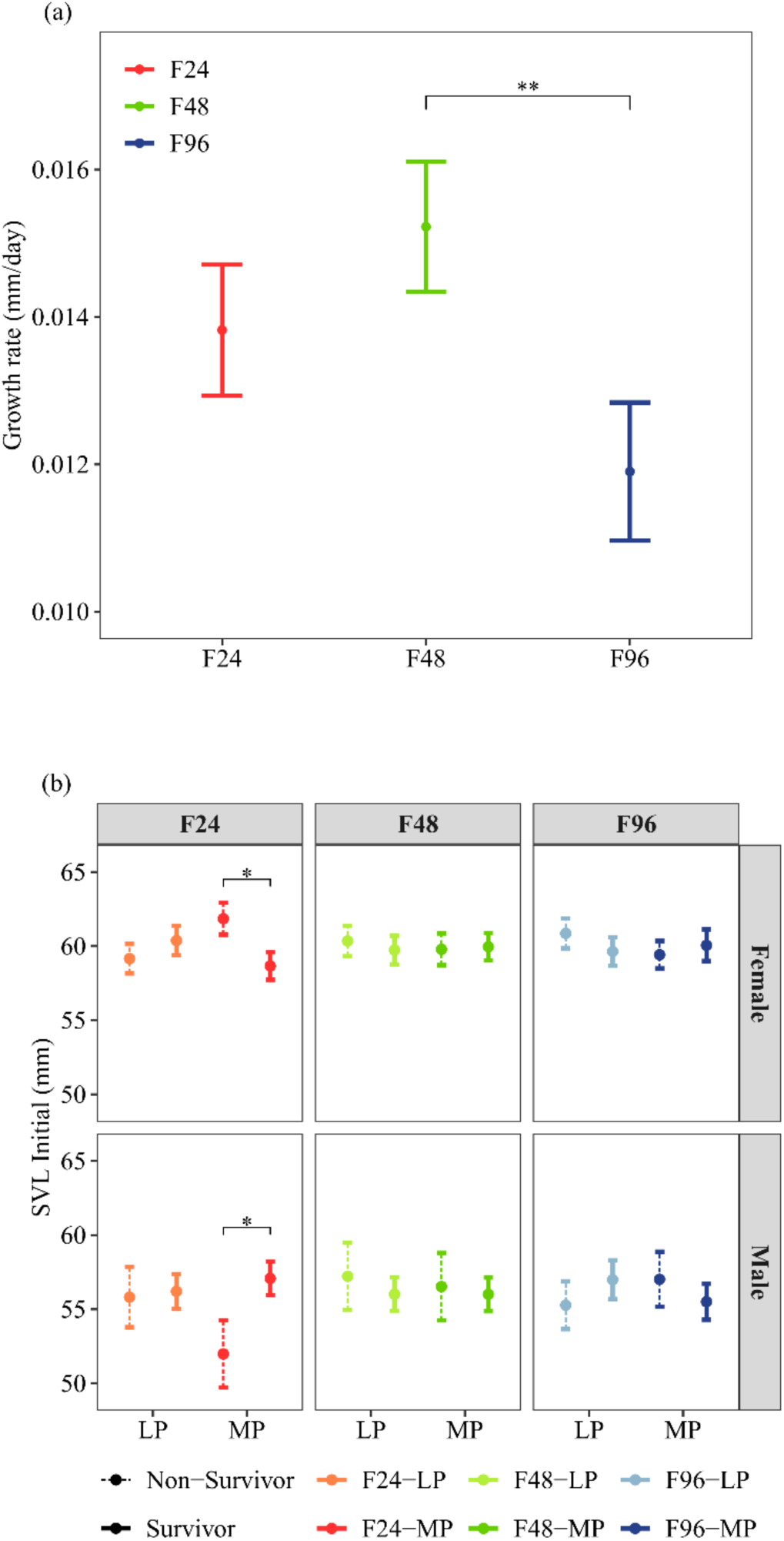
Differences among treatment levels in growth (**a**) and size-dependent survival of adults (**b**). Because the final body size of the non-survivors was unknown, the analyses were based on body size at release (SVL_Initial_). Predicted means ± SE for each combination of the frequency treatment (FREQ) (**a**) and the FREQ, PRED (predictability treatment), sex, and survival (**b**) are shown. FREQ levels are delimited as F24 (24-hour recurrence interval), F48 (48-hour interval), F96 (96-hour interval), and PRED levels as LP (less predictable) and MP (more predictable). Dashed lines in panel (**b**) correspond to non-surviving individuals, and solid lines represent survivors. Horizontal lines within graphs indicate significant post-hoc contrasts: * *P* < 0.05; ** *P* < 0.01; *** *P* < 0.001.

### Yearlings

In yearlings, there were no treatment differences in SVL at release (all *P* > 0.5; supplementary Fig. S3a) and treatments had no significant effect on survival and SVL at the end of the experiment (all *P* > 0.2; supplementary Fig. S3b). However, there were significant interactions of FREQ, sex, and survival (*χ^2^*_2_ = 9.297, *P* = 0.010), and PRED, sex, and survival (*χ^2^*_2_ = 6.034, *P* = 0.014) on SVL_Initial_. The first interaction exists because, in F48, females that survived had larger initial body sizes compared to females that did not survive (*P* = 0.048; Fig. 3a), while smaller females survived in F96 (*P* = 0.006; Fig. 3a), and no other significant contrasts existed. Yearling growth rate was significantly influenced by an interaction between FREQ, SVL_Initial,_ and sex (*χ^2^*_2_ = 15.781, *P* < 0.001). In females the size-dependent decline of the growth rate was slower in F24 compared to F48 and F96 (*lstrends* post-hoc contrast: *P* = 0.030 and *P* = 0.010, respectively; Fig. 3b), and there were no significant slope differences between F48 and F96 females (*lstrends* post-hoc contrast: *P* = 0.508). However, growth rate of F96 females was significantly higher than that of F48 females (note that the two lines do not cross and that the line of F96 was above the F48 line, post-hoc contrast: *P* = 0.011; Fig. 3b). Therefore, differences between frequency treatments in size-dependent survival (i.e., bigger females survived in F48 and smaller females survived in F96; Fig. 3a), were compensated by differences in size-dependent growth (i.e., surviving F48 females exhibited a significantly lower growth rate than surviving F96 females; Fig. 3b).

**Figure 3.**
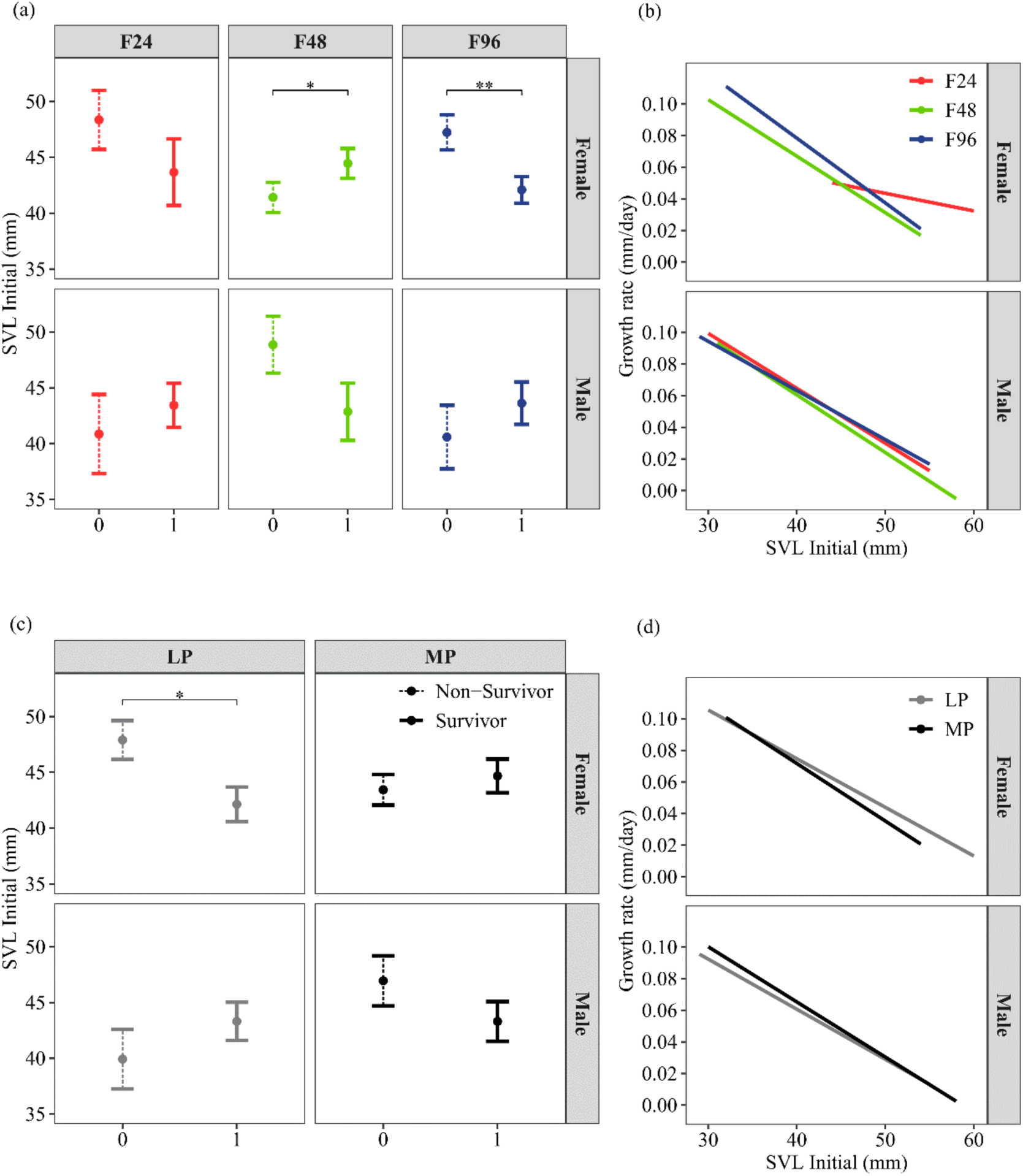
Treatment effects on body size-dependent survival (**a**, **c**) and growth rates (**b**, **d**) of yearlings. Predicted means ± SE for each interaction level of (**a**) FREQ (frequency treatment), sex, and survival and (**c**), PRED (predictability treatment), sex, and survival are shown, as well as the slopes between initial body size and growth rate for (**b**), the FREQ:sex interaction, and (**d**) the PRED:sex interaction. Abbreviations: F24, 24-hour recurrence interval; F48, 48-hour interval; F96, 96-hour interval; MP, more predictable; LP, less predictable. Dashed lines in panel (**a**) and (**c**) correspond to non-surviving individuals, and solid lines represent survivors. Horizontal lines indicate significant post-hoc contrasts: * *P* < 0.05; ** *P* < 0.01; *** *P* < 0.001.

In addition, growth rate was influenced by the interaction between FREQ, PRED, and sex (*χ^2^*_2_ = 15.437, *P* < 0.001), and SVL_Initial_ was highly significant (*F*_1,13_ = 106.190, *P* < 0.001). In MP, females in F48 grew significantly slower than those in F24 and F96 (*P* = 0.030 and *P* = 0.002, respectively; supplementary Fig. S4a), and no other contrasts were significant (all *P* > 0.1). In F48, females grew significantly faster in LP than MP (*P* = 0.008: supplementary Fig. S4a).

The second interaction exists because, in the less predictable treatment, surviving females were significantly smaller than non-survivors (*P* = 0.011; Fig. 3c), while no significant size differences between survivors and non-survivors existed in MP females and males. Growth rate was affected by an interaction between PRED, SVL_Initial,_ and sex (*χ^2^*_1_ = 9.900, *P* = 0.002, Fig. 3d). Neither in females nor in males did the slopes significantly differ between LP and MP treatments (all *P* > 0.4), but in females of the LP treatment, the bigger survivors grew more compared to the bigger survivors of the MP treatment. Differential growth thus compensated size-dependent survival, leading to no significant PRED treatment differences in final SVL (all *P* > 0.4; supplementary Fig. S3b).

### Juveniles

In juveniles, neither survival, growth, nor final SVL differed among treatments (all *P* > 0.2).

## Discussion

Current climate change leads to an increase in global temperatures, changes in the amount, variance, predictability, and frequency of precipitation^4,30,31^, and many other related phenomena. Climate change encompasses multiple dimensions and may involve complex interactive effects^18,19^. For example, less predictable climatic conditions are often associated with longer recurrence intervals (here mentioned as frequencies) of the studied climatic event^32^ and correlated changes have a greater impact than non-correlated changes do. Robust experimental proof for the impact of the joint action of different climatic parameters is limited, but essential for improving predictions of the effects of climate change on individuals, populations, and ecosystems.

Here, we used an experimental approach to examine the effects of precipitation frequency (FREQ) and precipitation predictability (PRED) on life-history traits of a short-lived lizard, the European common lizard (*Zootoca vivipara*). Our results show that FREQ, but not PRED, affected survival and growth of adult lizards (Fig. 1a, 2a). FREQ significantly affected adult survival in the 2019-20 experiment, but not in the 2020-21 experiment, and in males, F96 was the treatment with the lowest survival (Fig. 1a). In the 2019-20 experiment, temperature was significantly higher during the mating season in spring (March-May, Appendix S2: supplementary Figs. S5, S6) and the experimental populations received significantly more natural precipitation in May, July, and August, i.e., the main growth period of *Zootoca vivipara*, compared to the 2020-21 experiment (supplementary Figs. S5, S7). At first glance, these findings were unexpected because we predicted that longer frequencies (F96 treatment) would have less impact on survival in a year with more compared to a year with less overall precipitation.

Increased precipitation from May to August may have resulted from a greater number of precipitation events or alternatively from more precipitation per event. There were no significant differences between experimental years in the number of days with rain and the frequency of rainfall events (*X*^2^_1_ = 1.883, *P* = 0.170 and *X*^2^_1_ = 1.524, *P* = 0.375, respectively). Thus, treatment effects in 2019-20 on adult male survival cannot be explained by the recurrence interval of precipitation. This suggests that mainly the significantly higher spring temperature in the 2019-20 experiment explains the treatment-differences among years, suggesting that more precipitation per event could not offset the effect of higher temperatures. Higher ambient temperatures lead to higher water loss by means of transpiration^33^, and longer intervals between precipitation events in F96 compared with F24 and F48 reduce ambient humidity (independent of the higher precipitation per event), thereby increasing the risk of dehydration and mortality^34,35^. Vulnerability to higher temperatures and longer intervals of precipitation is particularly pronounced in ectotherms, as their performance, including reproduction, is highest at optimal body temperatures^36^. Furthermore, previous experimental evidence in *Z. vivipara* has shown that warmer temperatures are associated with an acceleration of the pace of life, leading to lower survival, higher metabolic costs, and restricted activity periods, which are particularly disadvantageous for older individuals^22^. In contrast to adult males, adult females had lower survival at higher frequencies (i.e., F24; Fig. 1a). These sex differences may exist due to the reproductive activities of the females (e.g., egg production that requires longer periods of thermoregulation), activity changes of adult females during the reproductive period^37^, and changes in the resolution of the conflict between water-saving responses and thermoregulatory behavior. Experimental results show that adult females living in drier habitats change their activity behavior to the early morning hours, when the habitat is cooler and wetter, while adult males living in drier habitats reduce their average daily activity^33^. In our experiment, supplemental precipitation events occurred early in the morning (before lizard activity started), and in F24, supplemental precipitation happened every day. Consequently, habitat humidity in F24 was higher each morning and temperature slightly cooler (compared to F48 and F96). Thus, on average adult F24 females may have become active later in the morning than adult F48 and F96 females, and the overlap with the adult males’ activity window may have been greater, potentially exposing females to more male sexual harassment and associated mortality^38^.

Conversely, in F48 and F96, adult females may have been active earlier in the morning every second day and on three of four days, due to the warmer and slightly drier soil (i.e., on the days without supplemental precipitation), and thus they may have suffered less from sexual harassment and associated death. Post-hoc analyses of mating scars present on the adult femalés belly (see supplementary Appendix S3, Fig. S8) are in line with this prediction, and frequency-dependent sexual harassment may explain why adult females in F24 survived worst. These results cannot be explained by life-history trade-offs (which involves trade-offs between survival and growth), because the simulated small changes in FREQ limited both survival and growth at the same time. Moreover, the results show that the responses to environmental changes are sex-specific^10,11^.

Adult SVL at the end of the experiment was affected by a significant triple interaction between FREQ, PRED, and sex, with some, but not all, combinations of this interaction being statistically significant (Fig. 1b). Observed differences between treatments are not due to the magnification of small non-significant differences in SVL_Initial_ (i.e., SVL at release) of adults (supplementary Fig. S2b), because at the beginning of the experiment, all treatments and enclosures exhibited very similar body sizes (i.e., mean and variance) and the small non-significant differences between treatments were not in the same direction as the significant differences in Fig. 1b.

Therefore, any differences observed at the end of the experiment arose either due to different growth rates (prediction 3) or differences in size-dependent survival (prediction 4). There existed a significant FREQ effect on adult growth rate (Fig. 2a), and the FREQ, PRED, and sex interaction was not significant (supplementary Fig. S2d), indicating that the significant triple interaction on SVL_Final_ of adults did not arise due to growth differences. There existed significant differences between adult survivors and non-survivors in SVL_Initial_, that depended on FREQ, PRED, and sex (Fig. 2b; Supplementary Appendix S1). The effects identified in this interaction align with those observed in SVL_Final_, leading to body size differences in adults at the end of the experiment (Fig. 1b; supplementary Figs. S2a, S2c). Consequently, in female and male adults, significant differences in SVL_Final_ are due to size-dependent survival (prediction 4) and are not explained by differences in growth rates. The direction of the predictability treatment on SVL_Final_ differed among the sexes (Fig. 1b), pointing to sex-specific strategies in response to precipitation predictability and precipitation frequency^7^, which is in line with earlier experimental findings^33^. In reptiles, body size varies significantly in relation to humidity and temperature^16,29^ and under favorable conditions, body sizes are bigger, reproductive success is greater^25^, and mate choice^24^ and phenology are affected by both parameters^23,39^. Female body size was largest in the F48-MP and F24-LP combinations (Fig. 1b), indicating that both combinations are favorable. Moreover, adult growth (Fig. 2a) and survival in 2019-20 (Fig. 1a) were highest in F48, suggesting that a precipitation frequency of 48 hours is the most suitable of the simulated regimes for adult common lizards. Surviving males exposed to F24-MP (and those exposed to F48-LP) were biggest (Fig. 1b), suggesting that they may have an advantage in inter- and intra-sexual interactions^40^ over those exposed to the other treatment combinations. In contrast, female body size was smaller in this treatment combination (Fig. 1b), suggesting that male body size also affects female body size and thereby female fecundity, and parturition date^38,41^, e.g., by means of sexual aggression. This suggests that the interaction between the frequency and predictability of precipitation has implications for sexual conflict and female reproductive success. More generally, these results show that precipitation predictability imposes costs on body size and survival (prediction 1) and that the direction of the precipitation predictability effect depends on precipitation frequency and sex. Thus, the precipitation predictability effects were more complex than predicted (prediction 1).

Reduced adult growth in the F96 treatment (Fig. 2a) and reduced survival of adult F96 males (Fig. 1a) are in line with prediction 1 and the idea that less frequent but larger precipitation events affect soil surface humidity^13,14^. It is also in line with studies showing that lower soil humidity leads to reduced growth rates of common lizards^16^, probably due to lower food availability^42^.

In yearlings, FREQ treatment significantly affected the size-dependent survival of females, but not that of males (Fig. 3a). Our results suggest that under shorter precipitation intervals (F48) food availability may have been higher^42^ favoring the survival of bigger females, while under lower food availability (F96), smaller females may have had an advantage. These results are consistent with previous findings showing that precipitation differences affect yearling females rather than yearling males^21^ and they may have arisen due to the greater energy requirements of females compared to males, i.e., because of the females’ larger body sizes^43^, and higher growth rates^21^. Body size-dependent survival of yearlings was also affected by an interaction between PRED and sex (Fig. 3c), suggesting again that males and females may have different strategies^7^. However, no significant effects of FREQ or PRED existed on yearling body size at the end of the experiment. This is because females showed faster growth rates in the treatments where survivors were smaller, compensating for the significant effects observed in size-dependent survival (Figs. 3a, 3c). This indicates that over the course of the experimental year, treatment differences in size-dependent survival of yearlings were compensated by differences in growth rates^44^ (predictions 2, 3, and 4). Smaller individuals may exhibit greater survival advantages under conditions of food scarcity, as their lower metabolic demands require less energy intake for maintenance^45^ and they may exhibit compensatory growth, whereby they accelerate their growth rate to offset the reduced body size (Figs. 3b, 3d)^44,46^. However, compensatory growth is predicted to be associated with higher mortality rates and reduced longevity^44,46^. Therefore, the fact that lower frequency and lower predictability favored the survival of smaller individuals over larger ones, with the survivors exhibiting catch-up growth, suggests detrimental effects on these individuals, consistent with life-history theory^10,11^.

In summary, we investigated how the frequency and predictability of precipitation influence life-history traits. The observed interactive effects of FREQ, PRED, age-class, and sex on size-dependent survival and SVL_Final_ were neither additive nor synergistic and led to treatment combination-specific effects^7,41^. Therefore, if each variable were examined independently, the results might not be generalizable. Considering frequency and predictability jointly is thus essential to understand responses to the predicted shifts in precipitation patterns and their effects on sexual selection, reproduction, and population dynamics. In addition, compensation mechanisms existed in yearlings, while adults were unable to compensate. Compensatory or catch-up growth has been observed across a wide range of species, including lizards, plants, invertebrates, and vertebrates^44,46^. Moreover, our results demonstrate that longer precipitation frequency negatively affects adult growth and survival, whereas predictability had no significant effects. This is similar to the findings of an earlier study^6^, which found no significant treatment-effects of precipitation predictability on survival when running simple survival analyses, but unraveled strong effects on population growth when using powerful seasonal age-structured matrix models. The here detected effects of precipitation frequency are in line with findings in nematodes, i.e., other ectotherms, whose population density decreased in presence of reduced precipitation frequency^47^. Moreover, in the dune grass *L. secalinus*, the percentage of seedling emergence was higher at higher precipitation frequencies^48^. In both studies, reduced precipitation frequency led to drought, whose consequences became increasingly studied in the last years.

Results originate particularly from plants^49^ and show that changes in precipitation frequency affect soil moisture^15^, vascular plant growth, and community dynamics^50^. In contrast, in this experiment, precipitation frequency was 24, 48, and 96 hours (1, 2, 4 days), and neither dry nor drought periods existed (i.e., lizards had 24 hours access to *ad libitum* water in the water ponds). Given the evidence for frequency effects in nematodes, plants, and reptiles, our results suggest that in a wide range of species the effects of small changes in precipitation frequency (even in the absence of water shortage) may importantly depend on precipitation predictability and that only considering both will unravel how they shape life-history traits and affect population dynamics of ectotherms, i.e., of the vast majority of terrestrial animals.

## Materials and methods

### Study species and semi-natural populations

*Zootoca vivipara* is a small ovoviviparous, hygrophilous lacertid (snout-to-vent length [SVL]: adult males: 40-61 mm, adult females: 46-76 mm) inhabiting humid habitats and high moisture substrates, including peat bogs and humid heathlands^51^. This species mainly inhabits boreo-alpine climates characterized by cold temperatures, moderate to high precipitation, and very humid habitats^52^. Three age classes can be distinguished based on body size and coloration^53^: juveniles (born in the current activity season), yearlings (born in the previous activity season), and adults (born ≥ 2 activity seasons ago). 12 independent semi-natural *Z. vivipara louislantzi* populations were established at the ecological field station of “El Boalar” (Instituto Pirenaico de Ecología, Jaca, Spain; 42°330 N, 0°370 W, 700 m a.s.l.; for details, see^6,21^ in mesocosms of 100 m^2^. Mesocosms contained standardized natural vegetation, hides, rocks, and water ponds, and were exposed to identical environmental conditions (light, temperature, natural precipitation), as they were located in the same meadow. Natural populations exhibit population densities ranging from 0.1 individuals/100m^2^ to 21.25 individuals/100m^2^ (^54,55^, and personal observations from the populations where the used lizards stem from). In natural populations in the central Pyrenees, precipitation occurs year-round in the form of rain, snow, mist, and dew, and precipitation exhibits no seasonal pattern (^56^ ; personal precipitation measures). At the field station, habitat humidity during the lizard’s activity period (March-October) is frequently below the lizards’ requirements and supplemental precipitation is required to provide them with the preferred humidity conditions.

### Precipitation treatments

We exposed mesocosms (hereafter referred to as enclosures) to different levels of precipitation frequency (FREQ) and intrinsic predictability (PRED) from March 15^th^ to October 15^th^ to wi This experiment was conducted in two consecutive experimental years (experimental year: June to May of the following year) to understand its generality. Frequency is defined as the average inter-occurrence time^57^. Three FREQ treatments were established, each in four enclosures: F24 (supplemental precipitation events occurred at 8 a.m. every 24 hours), F48 (every 48 hours), and F96 (every 96 hours). Total precipitation (i.e., supplemental plus natural precipitation) exhibited significant FREQ treatment differences (*χ^2^*_2_ = 30.875, *P* < 0.001; all pairwise contrasts: *P* < 0.05; Appendix S4, supplementary Fig. S9a). Additionally, two populations of each FREQ treatment were exposed to higher (MP: more predictable) and another two to lower (LP: less predictable) intrinsic predictability of precipitation. To this end, supplemental precipitation events happened at 8 a.m. in the MP treatment, whereas in the LP treatment, only half of the supplemental precipitation was provided at 8 a.m. and the other half was randomly (i.e., for every enclosure at different random times) provided throughout four consecutive days (supplementary Fig. S9b).

The experimental precipitation falling outside the lizard’s activity window simulated the daily morning dew falling every morning in the natural Pyrenean lizard populations from where the study lizards originated from (>1600 m a.s.l.). Differences in total precipitation (natural precipitation [see supplementary Appendix S2] and supplemental precipitation) among treatments were confirmed using weighted permutation entropy, calculated for each enclosure (for calculation details, see^58^). Permutation entropy is inversely related to intrinsic predictability, which is a measure of time series complexity^59^. Permutation entropy was larger in the LP treatment (0.94) and smaller in the MP treatment (0.75), showing that precipitation was less predictable in the LP treatment. Over four days, all enclosures received the same amount of precipitation (equivalent to 16 minutes of irrigation), which avoided drought events that may hinder the interpretation of the results due to statistical conflation.

In the Pyrenees and on the field site, lizard summer activity is bimodal, with little activity in the early morning (from 8 to 9 a.m.; all time specifications refer to winter time), a peak of morning activity between 10 and 12 a.m., and some activity in the afternoon (between 4 and 7 p.m.; personal observations). In the MP treatment, supplemental precipitation did not overlap with the peak of lizard activity, while in the LP treatment, it may have overlapped if the random precipitation was falling into the lizard’s activity window.

### Experimental conditions

Lizards were released in June 2019 and 2020. After hibernation, enclosures were searched for gravid females with big eggs, which were maintained in the laboratory until egg laying (for more details on female capture, laboratory protocol, egg incubation, and juvenile measurements, see^6,21^. At the end of the mating season (May/June), all alive lizards were captured by hand, brought to the laboratory for measurements, and kept in individual terraria under standardized conditions until release. Lizards were randomly released into the empty enclosures to initiate the second experimental year (for more details, see supplementary Appendix S5). In all enclosures, the same number of adults and yearlings was released (*N* = 20 in Exp. 2019-20 and *N* = 18 in Exp. 2020-21; for details see supplementary Table S4), and there were no significant differences between treatments in the number of released juveniles (*χ^2^*_1_ = 1, *P* = 0.549). There were also no significant treatment differences in body size (all *P* > 0.8), body condition (all *P* > 0.2), tail length (all *P* > 0.3), and frequency of adult male color morphs (all *P* > 0.7). All measures were taken blind with respect to the treatment/enclosure of origin. The capture, recapture, maintenance and handling of lizards was carried out under license granted by the Gobierno de Aragón (LC/ehv 24/2010/105 and 106) and with the approval of the used experimental protocols by the same organization (INAGA/500201/202071136). All procedures complied with current Spanish legislation and adhered to the ethical guidelines established by the Association for the Study of Animal Behavior the Animal Behavior Society for the use of animals in behavioral research, and the ARRIVE guidelines.

### Measures and statistical analysis

Recaptured lizards were measured for body size (accuracy: 1 mm) and body mass (accuracy: 0.001 g). Molt, mating scars, tail condition, male coloration, and female gravidity were recorded. Survival (yes/no) was determined at the end of the experimental year and corresponds to whether or not a released lizard was recaptured (note: during the following year, no unrecaptured lizards appeared). Growth rate (daily increase in SVL) was calculated by dividing SVL at recapture minus SVL at release by the number of days between release and recapture (days spent hibernating [between 1^st^ November and 1^st^ March] were not counted)^7^. Treatment effects on life-history traits were analyzed with mixed models from packages *nlme* and *lme4* in R (version 4.4.0). Linear mixed models with *Gaussian* error distribution were applied for body size, growth rate, and size-dependent survival, and generalized mixed models with *binomial* error distribution were fitted to analyze the probability of survival. Growth rate models included SVL_Initial_ as a covariate because larger individuals tend to exhibit a slower growth rate^60^.

Each age class was analyzed independently, and all initial models contained both treatments (i.e., FREQ and PRED), sex, and experimental year as factors, as well as their interactions. The combination of enclosure and year was modeled as a random effect (nested within treatment combination) in all presented analyses, as well as “Animal ID” for repeated measurements of adults, and “Mother ID” when analyzing juveniles, given that individuals from the same clutch are not independent. This approach avoids pseudoreplication by accounting for non-independence within the same experimental enclosures or repeated measures on the same lizards. Model selection was performed using backward elimination of non-significant terms^61^. Model assumptions (e.g., normality and heteroscedasticity of the residuals) were tested, and if not met, transformations were applied and heteroscedasticity was corrected using the *varIdent* function.

For significant factors with more than two levels, post-hoc tests were performed using the *emmeans* package. Natural climatic data was analyzed using general linear mixed models with *Gaussian* error distribution for average daily temperature and with *Poisson* error and a *log* link for daily precipitation.

## Acknowledgments

For field assistance, we thank Maria Urieta Lardiés and Guillém Masó Ferrerons.

## Author contributions

P. S. Fitze obtained the financial means. P. S. Fitze and R. Vicente Moreno designed the methodology. R. Vicente Moreno, M. Urieta Lardiés, and P. S. Fitze collected the data, and R. Vicente Moreno measured the different traits and analyzed the data under the supervision of P. S. Fitze. R. Vicente Moreno and P. S. Fitze drafted the manuscript.

## Data availability statement

Data will be published in open access (at https://digital.csic.es), once the article has been accepted, and the link to the data will be added to the final version of the manuscript.

