## supplementary for "Precipitation frequency and predictability interactively affect lizard life-history traits in absence of water shortage"

### Index of supplementary information

**Appendix S1**. Treatment effects on life-history traits.

- **Figure S1, S2, S3 and S4**. Treatment effects on lizard life-history traits.

**Appendix S2**. Natural climate.

- **Figure S5.** Monthly natural precipitation (in mm) and average monthly temperature (ºC).
- **Figure S6 and S7**. Predictions of the best-fitting model for average monthly temperature (ºC) and natural precipitation (mm).
- **Table S1 and S2.** Model selection for average monthly temperature (ºC) and natural precipitation (mm).

**Appendix S3**. Scars on female bodies produced by males during the initiation of copulation.

- **Table S3.** Number of mating scars counted on females´ bellies during the 2019-20 experimental year.
- **Figure S8.** Number of mating scars counted on females´ bellies during the 2019-20 experimental year.

**Appendix S4**. FREQ and PRED treatments.

- **Figure S9.** Example of the total daily precipitation (in mm).

**Appendix S5**. Details of the used release design.

- **Table S4.** Number of adults and yearlings, and number ± SE of juveniles released in each enclosure, in each experimental year.

#### Appendix S1: Treatment effects on life-history traits

There existed significant differences between survivors and non-survivors in $SVL_{Initial}$ of adults, that depended on FREQ, PRED, and sex (Supplementary Fig. S2c). In F24, larger MP females died and smaller MP females survived, while the opposite occurred in LP females, leading to the observed differences in $SVL_{Final}$ of adult females (Supplementary Figs. S2a, S2c). The opposite existed in F48 females: in LP, slightly smaller and in MP, slightly bigger females survived (Supplementary Fig. S2c), and although these differences were not significant, they explain the significant differences in $SVL_{Final}$ among LP and MP in F48 (Fig. 1b, supplementary Figs. S2a, S2c). In F24 males, there was a significant difference in $SVL_{Initial}$ between survivors and non-survivors (Supplementary Fig. S2c), with smaller males dying and larger males surviving in MP. This difference led to significant differences in $SVL_{Final}$ between the LP and MP treatments in F24 males (Supplementary Figs. S2a, S2c). Consequently, in female and male adults, significant differences in $SVL_{Final}$ are due to size-dependent survival (prediction 4) and are not explained by differences in growth rates (prediction 3).

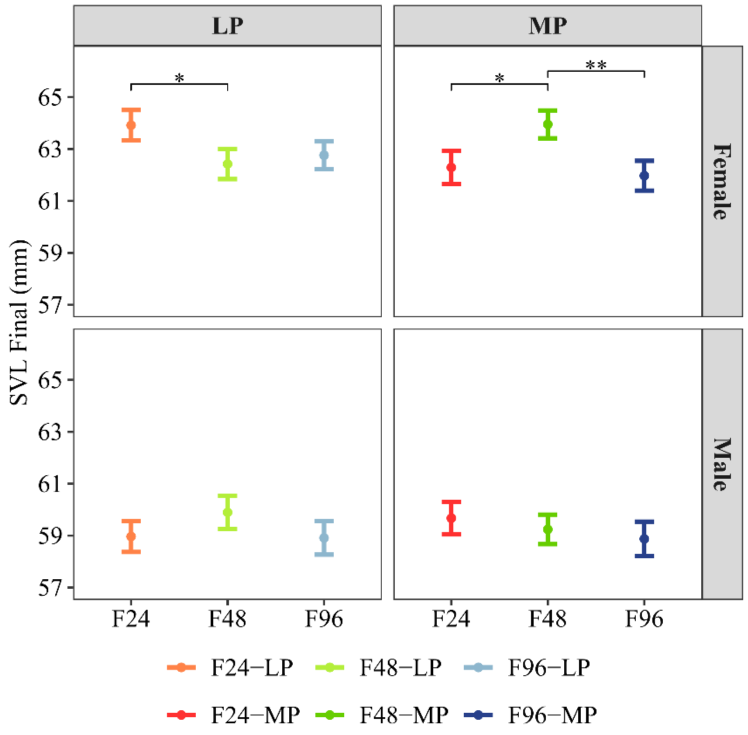


**Figure S1.** Treatment effects on adult body size at recapture. This graph shows the same values as Figure 1b, but the order of the treatment combinations is different to facilitate the understanding of the contrasts within the frequency (FREQ)-treatment. Predicted means ± SE per FREQ treatment (F24: 24-hour recurrence interval; F48: 48-hour interval; F96: 96-hour interval), predictability treatment (MP, more predictable; LP, less predictable), and sex are shown. Horizontal lines within graphs indicate significant post-hoc contrasts: * $P < 0.05$; ** $P < 0.01$; *** $P < 0.001$.

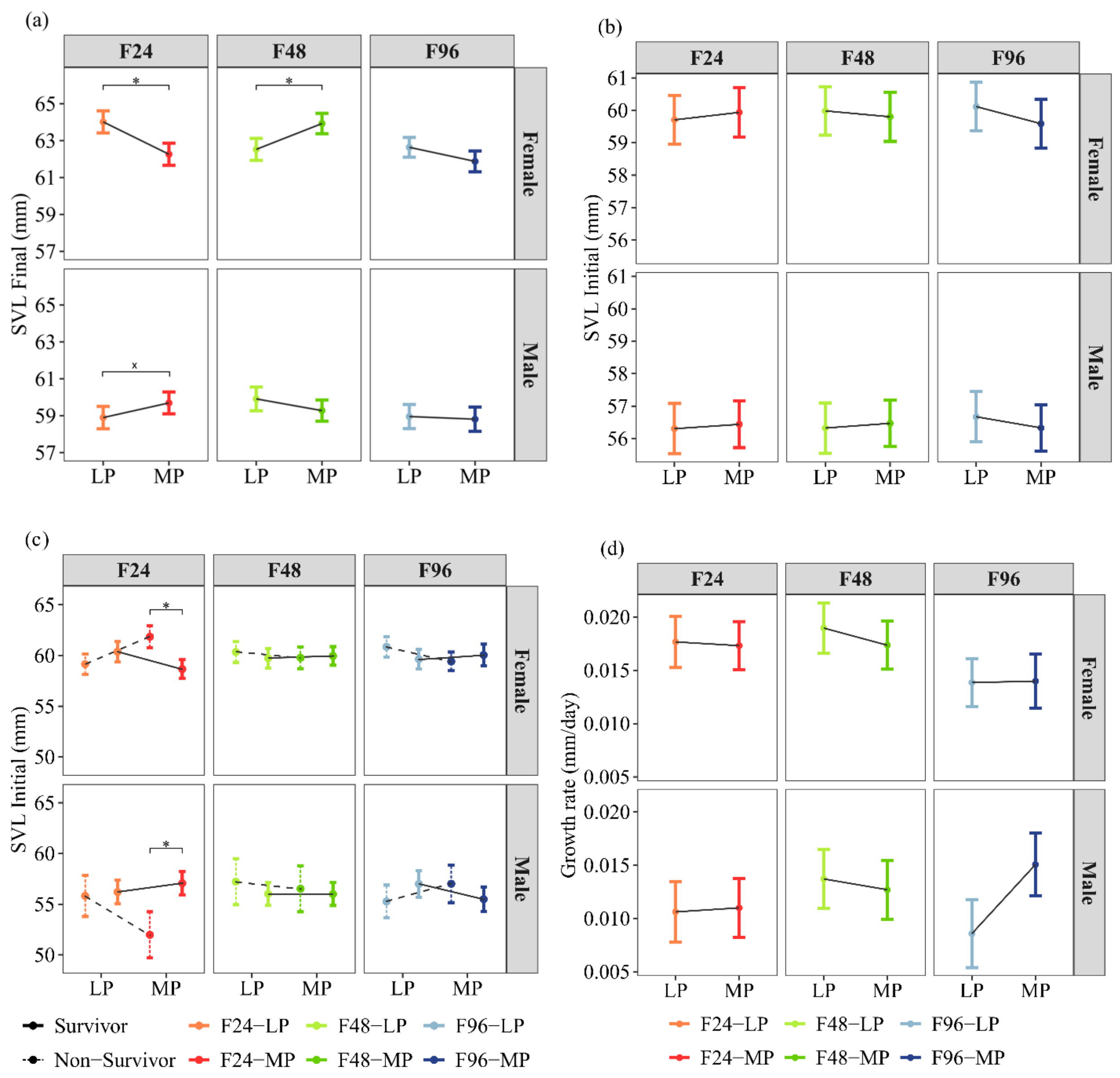


**Figure S2.** Summary overview facilitating the visual interpretation of the causes of the observed significant treatment effects on adult $SVL_{Final}$. (**a**) same figure as Figure 1b (FREQ, PRED, and sex interaction on adult $SVL_{Final}$). (**c**) same figure as Figure 2b (FREQ, PRED, sex, and survival interaction on adult $SVL_{Final}$). Added lines between treatment levels visualize the direction of the treatment effects. Interactions between FREQ, PRED, and sex on adult $SVL_{Initial}$ (i.e., at release) and adult growth rate were not significant and are shown in (**b**) and (**d**). Abbreviations: F24, 24-hour recurrence interval; F48, 48-hour interval; F96, 96-hour interval; MP, more predictable; LP, less predictable. Horizontal lines above means ± SE indicate significant post-hoc contrasts: * $P < 0.05$; ** $P < 0.01$; *** $P < 0.001$.

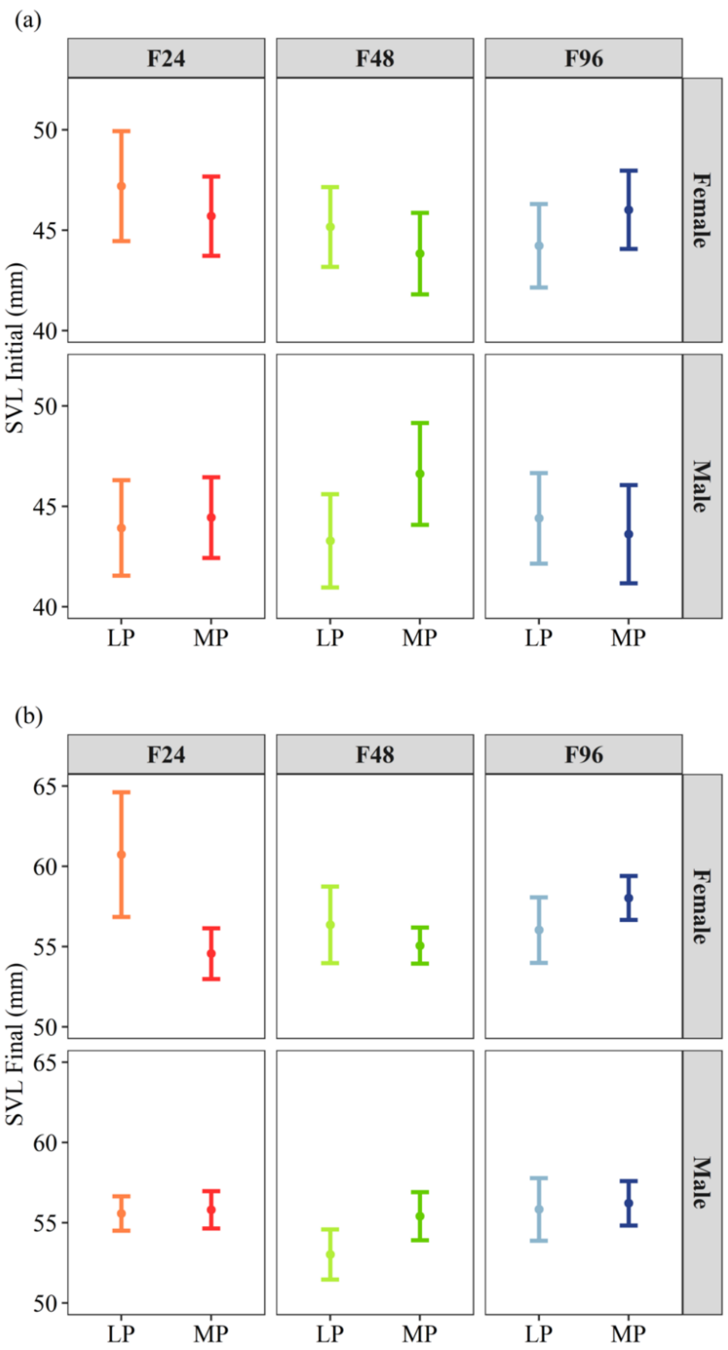


**Figure S3.** Interactions between FREQ, PRED, and sex on (**a**) yearling $SVL_{Initial}$ (i.e., at release) and (**b**) yearling $SVL_{Final}$ were not significant. Abbreviations: F24, 24-hour recurrence interval; F48, 48-hour interval; F96, 96-hour interval; MP, more predictable; LP, less predictable. Horizontal lines above means ± SE indicate significant post-hoc contrasts: * $P < 0.05$; ** $P < 0.01$; *** $P < 0.001$.

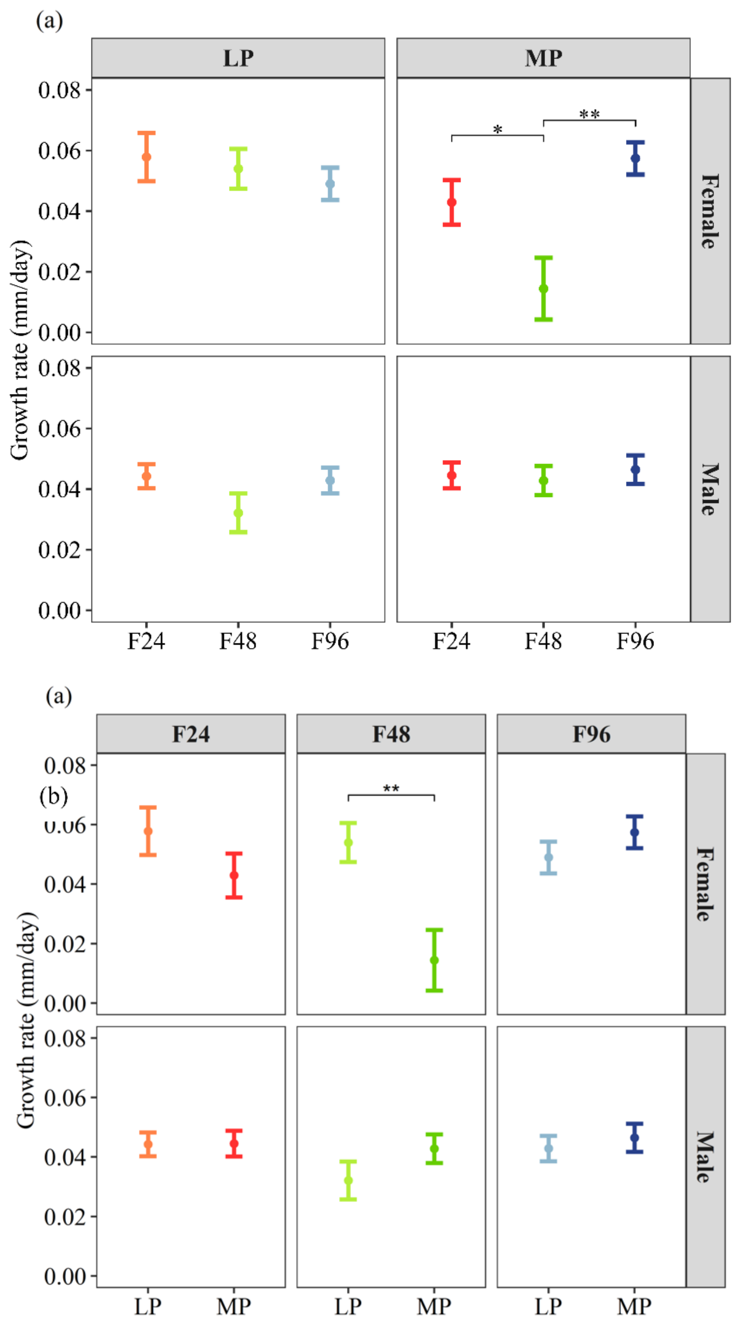


**Figure S4.** Treatment effects on yearling growth rate. Panel (**a**) shows the same values as panel (**b**), but the order of the treatments is different to facilitate the understanding of (**a**) the within FREQ and (b) the within PRED contrasts. Predicted means ± SE per FREQ treatment (F24: 24-hour recurrence interval; F48: 48-hour interval; F96: 96-hour interval), PRED treatment (MP, more predictable; LP, less predictable), and sex are shown. Horizontal lines within graphs indicate significant post-hoc contrasts: * $P < 0.05$; ** $P < 0.01$; *** $P < 0.001$.

### Appendix S2: Natural Climate

Over the course of the two experimental years (June 2019 to May 2020 and June 2020 to May 2021), environmental predictability of precipitation was manipulated at the enclosure level. During this period, the average temperature and the amount of precipitation were recorded on an hourly basis by a measurement station located on the field site. Since the beginning of the experimental year, temperatures increased to their peak in July (21.7° ± 0.39 SE) and then decreased to their minimum in January (2.16° ± 0.49 SE; supplementary Appendix S2: Figure S5), after which they increased again until the end of the experimental year. Regarding precipitation, we observed that in February and September 2019-20 it rained the least, and in 2020-21 it rained the least in March, July, and August. In both experimental years in October, December, and January, i.e., when lizards were hibernating, it was raining similarly (supplementary Appendix S2: Figure S5). The highest monthly rainfall was recorded in November 2019-20, i.e., during the hibernation of the lizards. In this month, it rained more than twice as much as in the same month of 2020-21 and approximately one-third more than in December, i.e., the wettest month. In the 2019-20 experiment, February was dry and March wet, while in 2020-21, the pattern was reversed (i.e., February was wet, and March dry). During the mating period, rainfall was most abundant in May 2019-20, followed by June 2020-21, and April, while in May 20-21 and June 2019-20 it was raining less (supplementary Appendix S2: Figure S5).

To determine whether there were differences in temperature and precipitation among the experimental years, we started with the simplest model including a linear trend and its interaction with experimental year and then included higher-order terms (and their interaction with experimental year) until the function was able to represented the pattern of the connected monthly averages and thereafter, until higher-order terms did not significantly improve the model.

According to the observed trends for average monthly temperature (supplementary Appendix S2: Figure S6), at least a cubic model, and for average monthly rainfall (supplementary Appendix S2: Figure S7), at least a quartic model was required to represent the observed pattern. Model selection was based on the Akaike Information Criterion (AIC)[1]. The best-fitting model was selected for each polynomial, and if several best-fitting models existed, the model with the fewest parameters, i.e., with greater parsimony, was prioritized. A significant quartic relationship existed between month and average monthly temperatures, which was significantly better than the best cubic model ($\Delta$AIC = 227.6) and not significantly different from the quintic models ($\Delta$AIC $\leq$ 1.31; supplementary Appendix 2: Table S1). All parameters of the selected quartic model were significant (year: $F_{1,723} = 9.0511$, $P = 0.003$; month: $F_{1,723} = 303.267$, $P < 0.001$; month$^2$: $F_{1,723} = 384.154$, $P < 0.001$; month$^3$: $F_{1,723} = 337.036$, $P < 0.001$; month$^4$: $F_{1,723} = 260.720$, $P < 0.001$; month$^3$*year: $F_{1,723} = 6.791$, $P = 0.009$; month$^4$*year: $F_{1,723} = 9.007$, $P = 0.003$), explaining 79% of the variation (supplementary Appendix S2: Figure S6). Post-hoc tests performed using the *emmeans* package revealed significant differences between experimental years from March to May, i.e. from the emergence of hibernation to the mating season (post-hoc contrast: March, $P = 0.01$; April, $P < 0.001$; May, $P < 0.001$), and from June to September (Jun, $P = 0.003$; Jul, $P = 0.003$; Aug, $P < 0.004$; Sept, $P = 0.007$; Oct, $P = 0.02$), i.e. the main annual growth period[2].

Quintic models of monthly precipitation were significantly better than quartic and all lower level models ($\Delta$AIC $\geq$ 4.48; supplementary Appendix S2: Table S2). Thus, high-order terms were tested, and the best-fitting model was a septic model 1i2567, which was significantly better than the best sextic model ($\Delta$AIC=124.68), more parsimonious than the other septic models and the octic models (supplementary Appendix S2: Table S2). All parameters of the selected septic model, except the main effect of year, were significant (year: $X^2_1 = 1.158$, $P = 0.282$; month: $X^2_1$

$= 167.078$, $P < 0.001$; month$^2$: $X^2{}_1 = 198.916$, $P < 0.001$; month$^5$: $X^2{}_1 = 163.487$, $P < 0.001$; month$^6$: $X^2{}_1 = 144.100$, $P < 0.001$; month$^7$: $X^2{}_1 = 126.723$, $P < 0.001$; month*year: $X^2{}_1 = 10.050$, $P = 0.002$; supplementary Appendix S2: Figure S7), explaining 43% of the variation (supplementary Appendix S2: Figure S6). Post-hoc tests using the *emmeans* package revealed significant differences between experimental years from October to May, i.e., hibernation, emergence, and the mating season (post-hoc contrast: Oct, $P = 0.028$; Nov-May, $P < 0.001$).

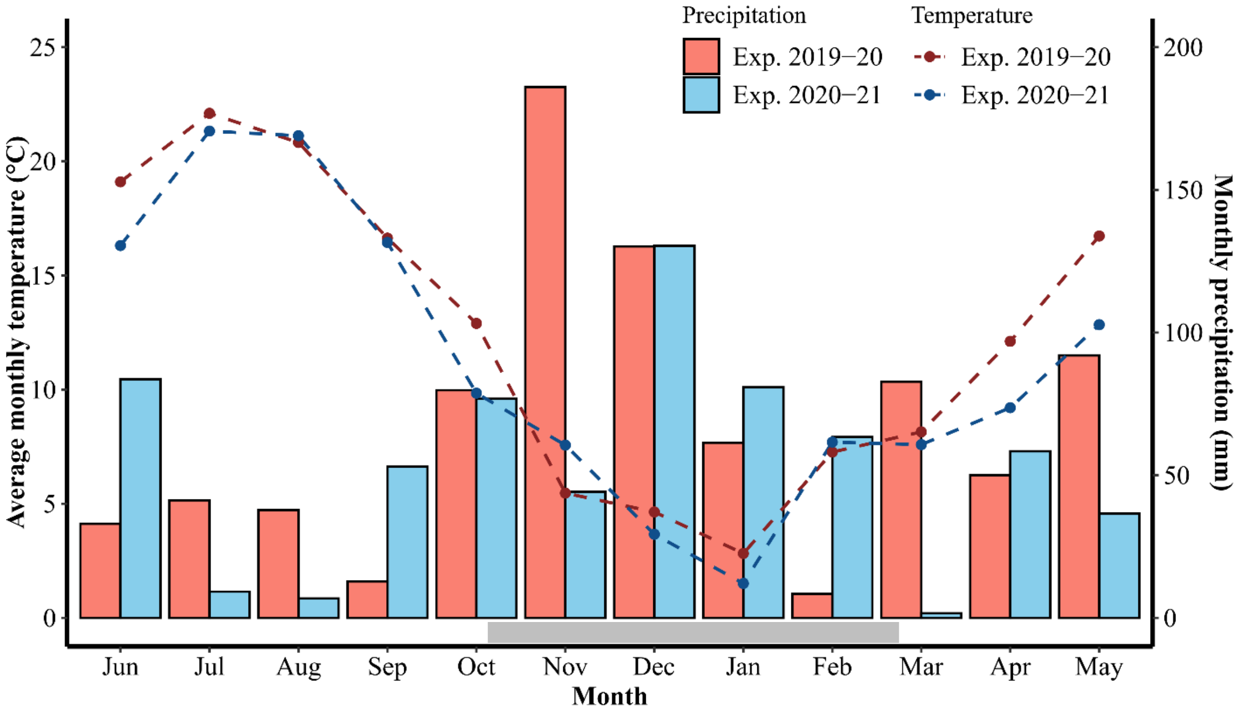


**Figure S5.** Monthly natural precipitation (in mm) and average monthly temperature (in °C) measured at the experimental field site over the duration of the experiment (2019-2020 and 2020-2021). Lines connect average monthly temperatures (red: experimental Year (Exp.) 2019.20; blue line: Exp. 2020-21). Bars correspond to monthly precipitation (red bars correspond to Exp. 2019.20; blue bars: Exp. 2020-21). The grey horizontal bar at the bottom of the precipitation bars corresponds to the months when lizards are hibernating. An experimental year began in June and ended in May of the following year

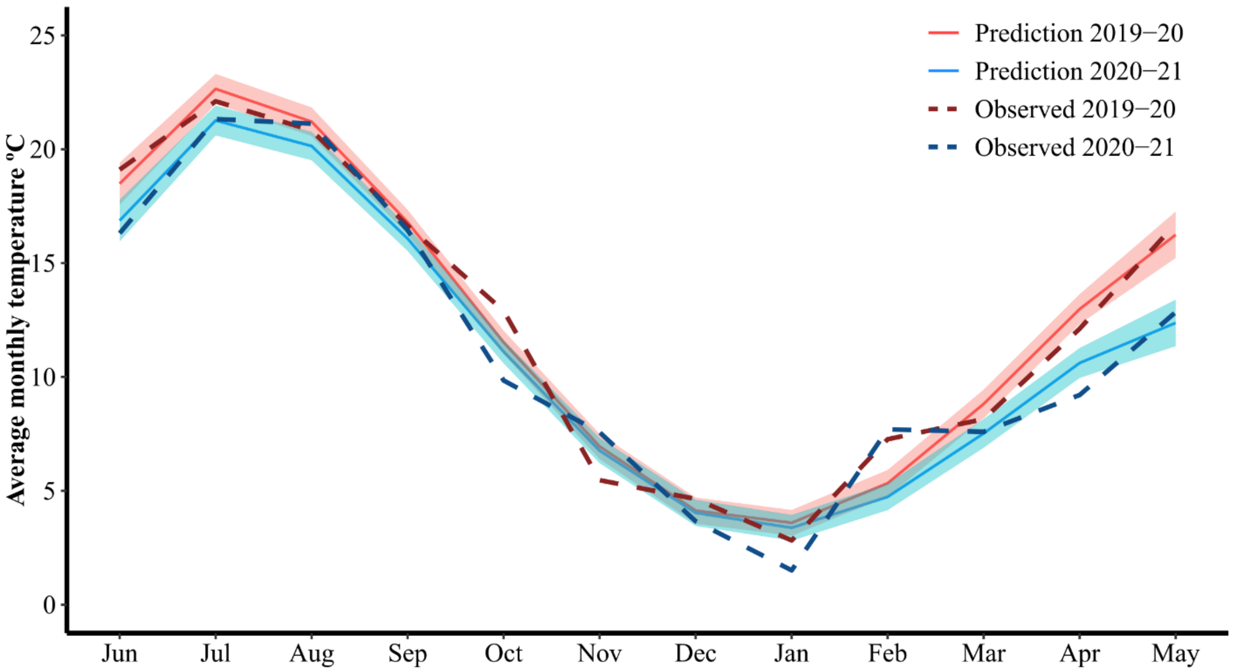


**Figure S6.** Predictions of the best-fitting model 123i4i (supplementary Appendix S2: Table S1) for average monthly temperature (°C) measured at the experimental field site over the course of the experiment (red line: years 2019–20; blue line: years. 2020–21). Each experimental year began in June and concluded in May of the following year.

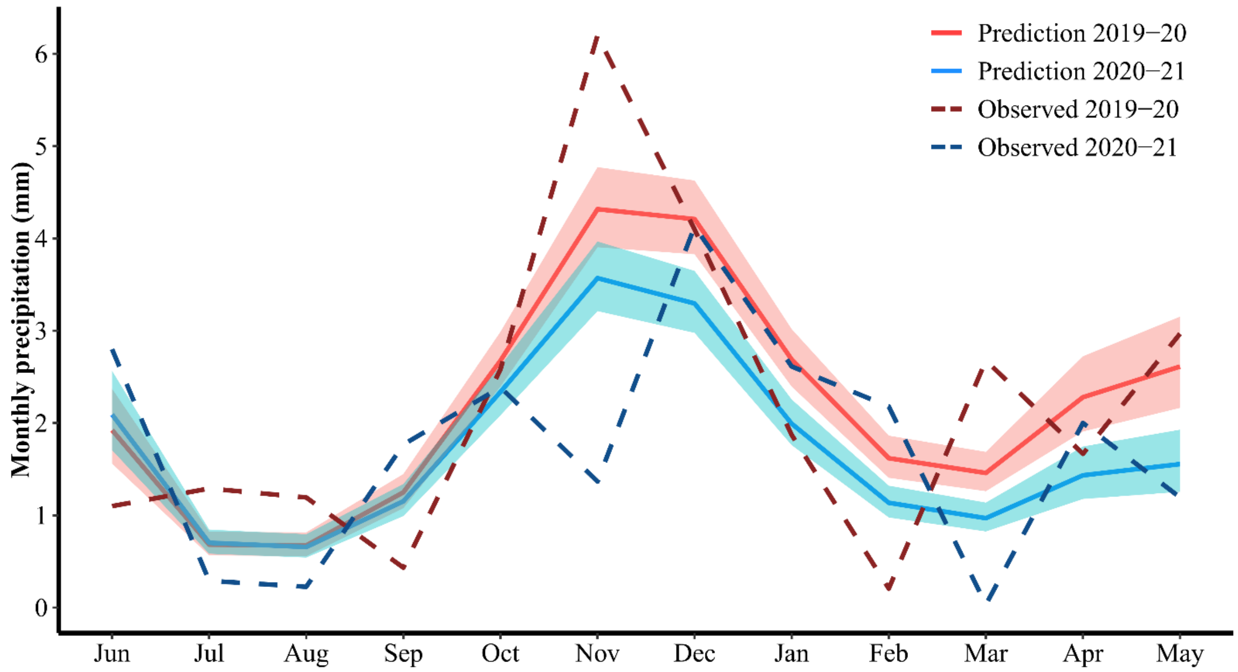


**Figure S7.** Predictions of the best-fitting model 1i2567 (supplementary Appendix S2: Table S2) for precipitation (mm) measured at the experimental field site over the course of the experiment (red line: years 2019-20; blue line: years 2020-21). Each experimental year began in June and concluded in May of the following year.

**Table S1. Model selection for average monthly Temperatures.**

Model selection started with the simplest model, including a linear trend and its interaction with year, followed by including the next higher-order term (and its interaction with year) until the function represented the pattern of the connected monthly averages (supplementary Appendix S2: Figure S5). Thereafter, higher-order terms (and their interaction with year) were included until they did not significantly improve the model. According to the observed trends (supplementary Appendix S2: Figure S5), average monthly temperature needs to be modeled with at least a cubic model. Model selection was based on the corrected Akaike Information Criterion (AICc), and a difference of $\geq 2$ AICc was considered to be significantly better[3]. In the case that several best-fitting models (BEST) existed, the more parsimonious (MP) model was prioritized (i.e. the model with the fewest parameters). Model abbreviations are as follows: numbers reflect power (i.e., 1 = linear, 2 = quadratic, 3 = cubic, etc), and 'i' behind the number indicates its interaction with year. The number (*N*) of parameters, AICc, ΔAIC, Aikaike Weights, and the used model selection criteria (BEST or MP). Selected models are bold.

| Model | N Parameter | AICc | ΔAIC | Weights | Selection Criteria |
|---|---|---|---|---|---|
| 1) linear models | | | | | |
| **1** | **4** | **4701.74** | **0.00** | **61.02** | BEST & MP |
| 1i | 5 | 4702.98 | 1.24 | 32.84 | |
| 0 | 3 | 4706.33 | 4.59 | 6.14 | |
| 0i | 3 | 4925.51 | 233.77 | 0.00 | |
| 2) quadratic models | | | | | |
| 12i | 6 | 4312.26 | 0.00 | 64.62 | |
| **12** | **5** | **4313.46** | **1.21** | **35.38** | MP |
| 1 | 4 | 4701.74 | 389.48 | 0.00 | |
| 3) cubic models | | | | | |
| **123i** | **7** | **4041.75** | **0.00** | **92.16** | BEST |
| 123 | 6 | 4046.67 | 4.93 | 7.84 | |
| 12 | 5 | 4313.46 | 271.72 | 0.0 | |
| 4) quartic models | | | | | |
| **123i4i** | **9** | **3814.15** | **0.00** | **97.07** | BEST |
| 123i4 | 8 | 3821.15 | 7.00 | 2.93 | |
| 123i | 7 | 4041.75 | 227.60 | 0.00 | |
| 5) quintic models | | | | | |
| 123i4i5 | 10 | 3813.45 | 0.00 | 48.3 | |
| **123i4i** | **9** | **3814.15** | **0.70** | **34.0** | MP |
| 123i4i5i | 11 | 3815.45 | 2.01 | 17.7 | |

**Table S2. Model selection for average monthly Precipitation.**

Model selection started with the simplest model, including a linear trend and its interaction with year, followed by including the next higher-order term (and its interaction with year) until the function represented the pattern of the connected monthly averages (supplementary Appendix S2: Figure S5). Thereafter, higher-order terms (and their interaction with year) were included until they did not significantly improve the model. According to the observed trends (supplementary Appendix S2: Figure S5), monthly precipitation needs to be modeled with at least a quartic model. Model selection was based on the corrected Akaike Information Criterion (AICc), and a difference of $\geq 2$ AICc was considered to be significantly[3]. In the case that several best-fitting models (BEST) existed, the most parsimonious (MP) model was prioritized (i.e. the model with the fewest parameters). Model abbreviations are as follows: numbers reflect power (i.e., 1 = linear, 2 = quadratic, 3 = cubic, etc) and 'i' behind the number indicates its interaction with year. The number (*N*) of parameters, AICc, ΔAIC, Aikaike Weights, and the used model selection criteria (BEST or MP). Selected models are bold.

| Model | N Parameter | AICc | ΔAIC | Weights | Selection Criteria |
|---|---|---|---|---|---|
| 1) linear models | | | | | |
| **1i** | **4** | **5512.35** | **0.00** | **95.09** | BEST |
| 1 | 3 | 5518.28 | 5.93 | 4.90 | |
| 0i | 2 | 5529.84 | 17.48 | 0.02 | |
| 0 | 2 | 5535.64 | 23.29 | 0.00 | |
| 2) quadratic models | | | | | |
| **1i2** | **5** | **5435.21** | **0.00** | **59.02** | BEST & MP |
| 1i2i | 6 | 5435.94 | 0.73 | 40.98 | |
| 1i | 4 | 5512.35 | 77.14 | 0.00 | |
| 3) cubic models | | | | | |
| **1i2** | **5** | **5435.21** | **0.00** | **66.53** | BEST & MP |
| 1i23 | 6 | 5437.21 | 1.99 | 24.59 | |
| 1i23i | 7 | 5439.24 | 4.03 | 8.88 | |
| 4) quartic models | | | | | |
| 1i24 | 6 | 5435.11 | 0.00 | 42.00 | |
| **1i2** | **5** | **5435.21** | **0.10** | **39.88** | MP |
| 1i24i | 7 | 5436.79 | 1.68 | 18.12 | |
| 5) quintic models | | | | | |
| **1i25** | **6** | **5429.78** | **0.00** | **59.31** | BEST & MP |
| 1i25i | 7 | 5430.73 | 0.96 | 36.78 | |
| 1i2 | 5 | 5435.21 | 5.44 | 3.91 | |
| 6) sextic models | | | | | |
| **1i256** | **7** | **5278.97** | **0.00** | **60.46** | BEST & MP |
| 1i256i | 8 | 5279.82 | 0.85 | 39.54 | |
| 1i25 | 6 | 5429.78 | 150.81 | 0.00 | |
| 7) septic models | | | | | |
| **1i2567** | **8** | **5154.29** | **0.00** | **52.23** | BEST & MP |
| 1i2567i | 9 | 5154.47 | 0.18 | 47.77 | |
| 1i256 | 7 | 5278.97 | 124.68 | 0.00 | |
| 8) octic models | | | | | |
| **1i2567** | **8** | **5154.29** | **0.00** | **40.33** | BEST & MP |
| 1i25678i | 10 | 5154.75 | 0.45 | 32.13 | |
| 1i25678 | 9 | 5155.05 | 0.76 | 27.54 | |

#### Appendix S3: Scars on female bodies produced by males during the initiation of copulation

In *Zootoca vivipara*, copulation attempts are quite violent[1,2,3], since males bite females on the posterior abdomen with their mouth and sometimes try to force copulation against the female's will[2]. Once females are immobilized, they try to twist their tail below that of the female in order to copulate. Thereby, they produce mating scars on the females that can be seen even after several weeks.

In F24 of our experiment, supplemental precipitation events occurred every morning before the lizard's activity started. Since female activity depends on the ambient temperature, females were probably active later due to lower ambient temperatures, resulting in a larger overlap between the activity windows of females and males. This may have exposed females to more male sexual harassment and associated mortality[3]. Conversely, in F48, females may have been active later in the morning on every second day (i.e., on the days with supplemental precipitation). Moreover, in F96 females may have been active later on every fourth day, exposing them least to male mating attempts. Therefore, we predicted that the number of mating attempts, and thus the number of mating scars[1,4], should follow the following order: F24 > F48 > F96. As visible in supplementary Appendix S3 Figure S8 and Table S3, F24 females exhibited the highest number of scars, followed by F48 and F96 females.

To test for statistical differences among frequency treatment, we used an ordered Heterogeneity (OH) test that accounts for the directional hypothesis[5]. The Spearman´s rank correlation ($R_s$) between the expected and observed order was 1, $r_s P_c$ was 0.81, and $P$ was $< 0.05$. This shows that the number of bites observed on the female's belly significantly depended on the frequency

(FREQ) treatment as predicted by treatment-induced differences in the daily onset of female activity. This points to FREQ-dependent harassment, which can explain why females in F24 exhibited the lowest survival.

**Table S3.** Number of mating scars counted on females´ bellies during the 2019-20 experimental year. Predicted means ± SE per FREQ (F24: 24-hour recurrence interval; F48: 48-hour interval; F96: 96-hour interval) and predictability (PRED) treatment (LP: less predictable; MP: more predictable) are shown.

| | F24 | F48 | F96 | Total |
|---|---|---|---|---|
| LP | 11.00 ± 2.19 | 11.50 ± 1.84 | 8.60 ± 2.39 | 10.47 ± 1.21 |
| MP | 14.75 ± 2.46 | 10.00 ± 1.85 | 8.00 ± 1.95 | 10.69 ± 1.25 |
| **Total** | 12.88 ± 1.66 | 10.81 ± 1.29 | 8.32 ± 1.52 | 10.57 ± 0.86 |

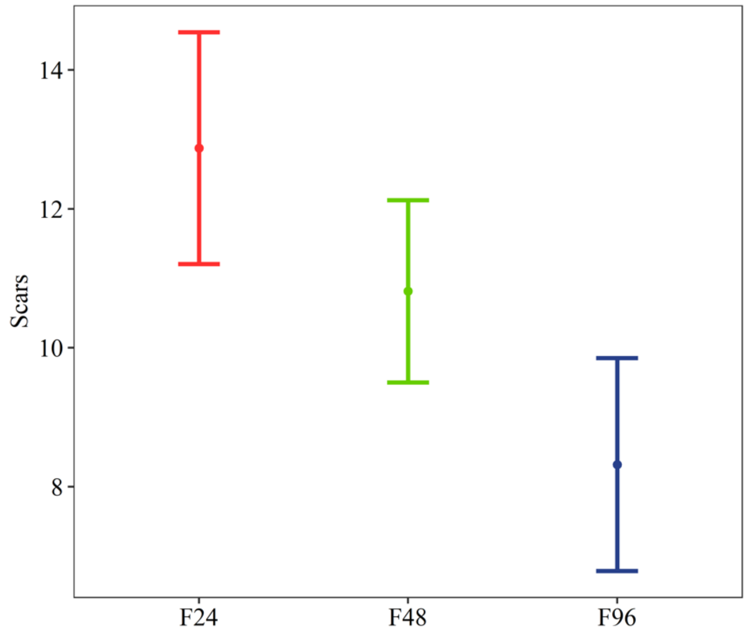


**Figure S8**. Number of mating scars counted on females´ bellies during the 2019-20. Predicted means ± SE per FREQ treatment are shown (F24: 24-hour recurrence interval; F48: 48-hour interval; F96: 96-hour interval).

### Appendix S4: FREQ and PRED treatments

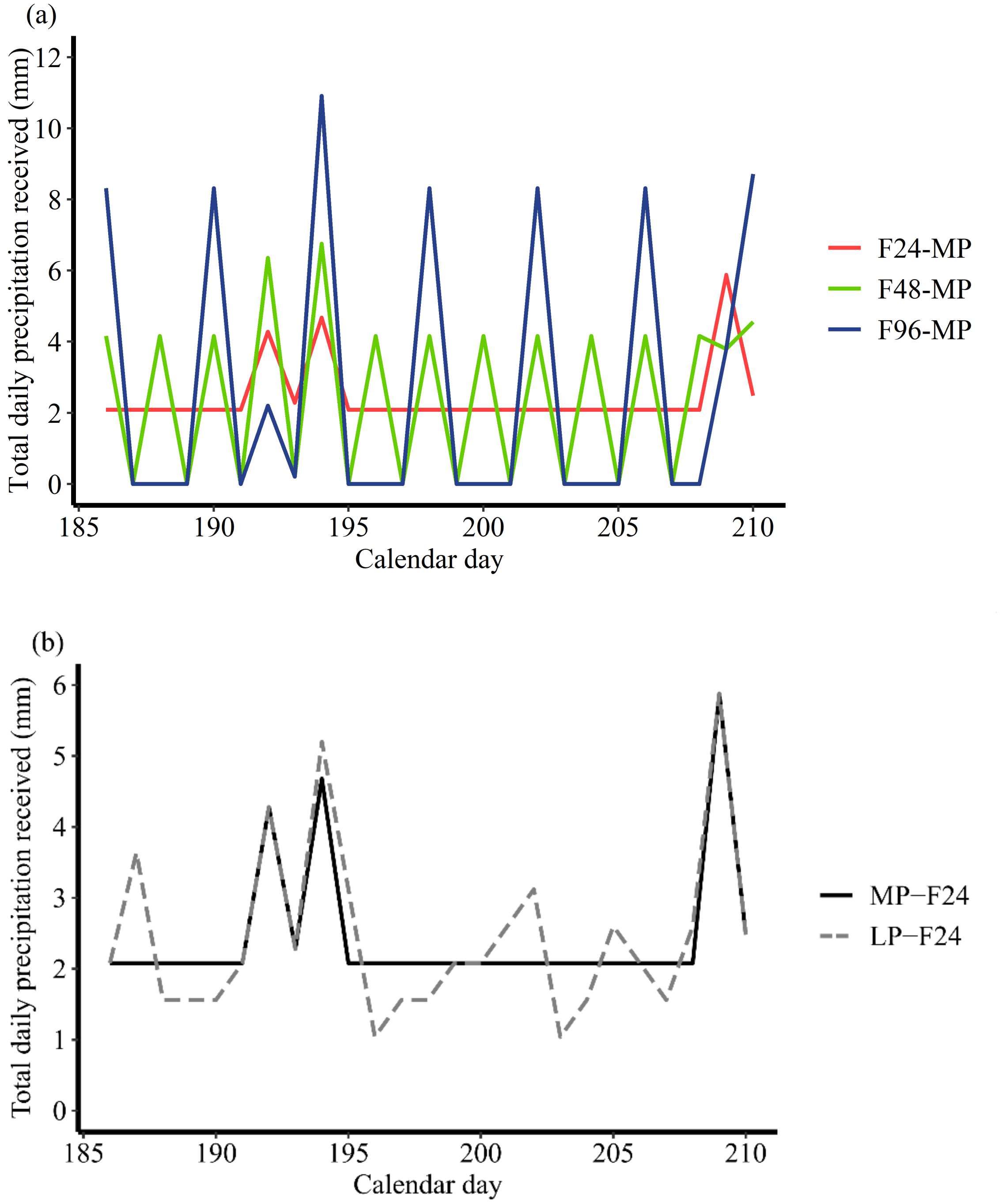


**Figure S9.** Example of the **total daily precipitation (in mm),** i.e., the sum of supplemental and natural precipitation, over a randomly selected period from June 4th to June 28th, 2020. **(a)** Total daily precipitation in the three levels of the frequency treatment (F24, 24-hour recurrence interval;

F48, 48-hour interval; F96, 96-hour interval) of the more predictable treatment (MP). (**b)** Total daily precipitation in the more (MP) and less predictable (LP) treatment of the F24 treatment. Supplemental precipitation events happened at 8 a.m. in the MP-F24 treatment on every day (see horizontal lines at 2.08 mm, on days without natural precipitation). In the LP-F24 treatment, half of the supplemental precipitation was as well provided on every day at 8 a.m. and the other half was randomly provided throughout four consecutive days. The average total daily precipitation was identical ($F < 0.001$, $P = 1.000$) across all treatment levels throughout the entire experiment (2019-2021).

#### Appendix S5: Details of the used release design

At the beginning of the experiment, lizards were randomly assigned to unfamiliar enclosures (i.e., they were not placed into the same enclosure where they were living previously). Individuals from previous treatments were evenly distributed among new treatments, and when possible, only one lizard of each previous enclosure was released into the same new enclosure. Hatchlings belonging to the same clutch were released in the same enclosure, not in the enclosure where their mother was living previously, nor in the enclosure where the mother was released, to avoid acclimation to a specific enclosure and mother-offspring competition[1]. After release, all enclosures contained the same number of adults and yearlings, and there were no differences between treatments in the number ($F_{2,18} = 0.600$, $P = 0.559$; supplementary Appendix S5: Table S4) and sex ratio ($F_{2,18} = 0.026$, $P = 0.975$; supplementary Appendix S5: Table S4) of the released juveniles. Adult sex ratio (32% males), was the same in each enclosure and similar to the sex ratio typically observed in natural populations (average: 39% males[2]).

**Table S4.** Number of adults and yearlings, and number ± SE of juveniles released in each enclosure, in each experimental year.

| | **Exp. 2019-20** | | | **Exp. 2020-21** | | |
|---|---|---|---|---|---|---|
| | *Adults* | *Yearlings* | *Juveniles* | *Adults* | *Yearlings* | *Juveniles* |
| *Female* | 11 | 1 | 4.50 ± 0.26 | 10 | 2 | 5.33 ± 0.22 |
| *Male* | 6 | 2 | 8.25 ± 0.28 | 4 | 2 | 7.92 ± 0.26 |
| ***Total*** | **17** | **3** | **12.75 ± 0.15** | **14** | **4** | **13.25 ± 0.30** |
